# Critical Thermal maximum measurements and its biological relevance: the case of ants

**DOI:** 10.1101/2020.12.09.417410

**Authors:** Chi-Man Leong, Toby P. N. Tsang, Benoit Guénard

**Affiliations:** School of Biological Sciences, The University of Hong Kong, Kadoorie Biological Sciences Building, Pok Fu Lam Road, Hong Kong SAR, China

**Keywords:** Methodology, Physiology, Upper thermal limit, Thermal ecology, Ants

## Abstract

1. Upper thermal limit (UTL) is a key trait in evaluating ectotherm fitness. Critical Thermal maximum CT_max_, often used to characterize the UTL of an organism in laboratory setting, needs to be accurate to characterize this significant and field-relevant threshold. The lack of standardization in CT_max_ assays has, however, introduce methodological problems in its measurement and incorrect estimation of species upper thermal limit; with potential major implications on the use of CT_max_ in forecasting community dynamics under climate change. In this study we ask if a satisfactory ramping rate can be identified to produce accurate measures of CT_max_ for multiple species.
2. We first identified the most commonly used ramping rates (i.e. 0.2, 0.5 and 1.0 °Cmin^−1^) based on a literature review, and determined the ramping rate effects on CT_max_ value measurements in 27 ant species (7 arboreal, 16 ground, 4 subterranean species) from eight subfamilies using both dynamic and static assays. In addition, we used field observations on multiple species foraging activity in function of ground temperatures to identify the most biologically relevant CT_max_ value to ultimately develop a standardized methodological approach.
3. Integrating dynamic and static assays provided a powerful approach to identify a suitable ramping rate for the measurements of CT_max_ values in ants. Our results also showed that among the values tested the ramping rate of 1 °Cmin^−1^ is optimal, with convergent evidences from CT_max_ values measured in laboratory and from foraging thermal maximum measured in the field. Finally, we illustrate how methodological bias in terms of physiological trait measurements can also affect the detection of phylogenetic signal (Pagel’s *λ* and Bloomberg’s *K*) in subsequent analyses.
4. Overall, this study presents a methodological framework allowing the identification of suitable and standardized ramping rates for the measurement of ant CT_max_, which may be used for other ectotherms. Particular attention should be given to CT_max_ values retrieved from less suitable ramping rate, and the potential biases that functional trait based research may induce on topics such as global warming, habitat conversion or their impacts on analytical interpretations on phylogenetic conservatism.

## 1. INTRODUCTION

Organisms are increasingly exposed to novel environmental conditions in responses to global changes such as deforestation, urbanization and climate change. High temperatures, in particular, limit species survival, reproduction and foraging activities; with ectotherms, can be limited by high temperatures (Deutsch *et al.*, 2008). Therefore, measuring ectotherms’ upper thermal limit is paramount to obtain valuable information to forecast changes in species diversity and distribution in response of increasing temperatures (Hoffmann, Chown & Clusella-Trullas, 2013).

Thermal performance curves represent useful indicators of an organism’s performance in function of the temperature experienced (Sinclair *et al.*, 2016); and delimit the range at which organisms remains active. The upper thermal maximum, referred as Critical Thermal maximum (CT_max_), is a particularly important threshold representing the temperature at which an organism is unable to withstand with heat stress (Lutterschmidt & Hutchison, 1997b). Investigating heat tolerance is paramount to study species fitness under climate change (Kellermann *et al.*, 2012b), however, forecasting the impacts of temperature increase is challenging due to the lack of standardized methods for CT_max_ measurement (Terblanche *et al.*, 2007). For instance, the ramping rates used (increase of temperature over time) to measure CT_max_ differ widely and produce major differences in CT_max_ values for a same species (Lutterschmidt & Hutchison, 1997b; Terblanche *et al.*, 2007; Chown *et al.*, 2009). Therefore, a comparable and biologically relevant method for CT_max_ measurement directly applicable in ecology is strongly needed to provide accurate values representing species’ maximal thermal limit.

CT_max_ was defined by Cowles and Bogert (1944) as *“the thermal point at which locomotory activity becomes disorganized and the animal loses its ability to escape from conditions that will promptly lead to its death”.* CT_max_ measurement is thus an experimental approach to reach the upper thermal limit point by increasing the environmental temperature until the loss of muscle control or heat-coma. The heat injury caused by ramping rate depends on two important dose-variables, the intensity of heat and the exposure duration (Jørgensen, Malte & Overgaard, 2019), with the use of different ramping rates resulting in variations of both variables (Terblanche *et al.*, 2007).

The use of particular ramping rate has been debated since CT_max_ implementation, with a tradeoff existing between the use of slow or fast ramping rates. Longer duration of experimental measurement in slow ramping rates affects CT_max_ values retrieved due to heat acclimation, a characteristic of phenotypic plasticity, and provides less accurate CT_max_ values (Cox, 1974; Hutchison, 1976; Hutchison & Maness, 1979). Moreover, the accumulation of heat injury is mainly dependent on exposure time rather than heat intensity within experiments using slow ramping rates (Chown *et al.*, 2009; Nyamukondiwa & Terblanche, 2010). On the other hand, fast ramping rates allow individuals to reach their upper thermal limit in a short period of time and with little heat injury caused by exposure duration. Besides, fast ramping rate can accurately measure CT_max_, as the individual body temperature can track the experimental temperature used (Barker, Townsend & Hacunda, 1981). An ideal velocity of ramping rate for CT_max_ should represent upper thermal limit for a given species to cease vital functions in the field (Lutterschmidt & Hutchison, 1997b). For instance, investigating CT_max_ for a given species can, in theory, forecast the temperature at which this species stops to forage as the Foraging Temperature maximum (FT_max_). Furthermore, in order to compare different studies, the satisfactory ramping rate should be standardized (Krebs, 1989), to provide a consistent method for a given taxon or across multiple taxa (Lutterschmidt & Hutchison, 1997b; Terblanche *et al.*, 2007). This lack of standardization, however, leaves ultimately the following question unresolved: *what is the biologically accurate CT_max_ value of a given species?*

In the present study, we used dynamic and static assays in ants, a model organism in ectotherms ecophysiology studies (Schumacher & Whitford, 1974; Lutterschmidt & Hutchison, 1997b; Kaspari *et al.*, 2015), to investigate the experimental measurement of CT_max_. Overall, our goal is to provide an overview of the limitations arising from the use of different methodologies and identify a more suitable protocol to measure biologically relevant CT_max_ values. To provide a general model for studying upper thermal limit in different taxa, we investigate the upper thermal limit in 27 ant species associated with different micro- and habitats, body sizes, phylogenetic clades, and biogeographic origins. First, we reviewed the different methodologies used to measure upper thermal limit in ants to show the breadth of methods used and select particular values to be tested in our following experiments. Second, we investigated how the intensity of heat and exposure duration influence upper thermal tolerance at both intra- and interspecific levels (Fig. 1 a & b) by using a mixed approach combining dynamic and static assays. Third, within an ant assemblage of 27 ant species, we predicted strong positive correlations in dynamic assays between the ramping rates used (°Cmin^−1^) and the CT_max_ values (°C) retrieved at both intraspecific (Q1 in Fig.1A) and interspecific levels (Q2 in Fig.1A) independently of the ecology, morphology or evolutionary history of the species. Fourth, in static assays we hypothesized a strong negative correlation between durations (minutes) and the ramping rate (°Cmin^−1^) used to retrieve the CT_max_ values (°C) (Q3 in Fig.1A). Moreover, as species show different tolerances of exposure duration in withstanding the temperature set for each species CT_max_ value (see Q3); faster ramping rates are predicted to decrease interspecific variation of exposure duration forecasted based on thermal tolerance landscapes (Rezende, Castañeda & Santos, 2014) (Q4 in Fig.1A). Fifth, we compared the CT_max_ values retrieved from dynamic assays obtained from three ramping rate treatments with the maximum activity temperatures observed (FT_max_) in year-long field observations (Fig.1B) to determine the concordance between field and laboratory approaches (Cerdá, Retana & Cros, 1998). Finally, we investigated whether CT_max_ is different between species with varying habitat and strata preferences, as well as if it is phylogenetically conserved, using CT_max_ measured by different ramping rates to assess if methodological differences can affect the findings.

**Figure 1A:**
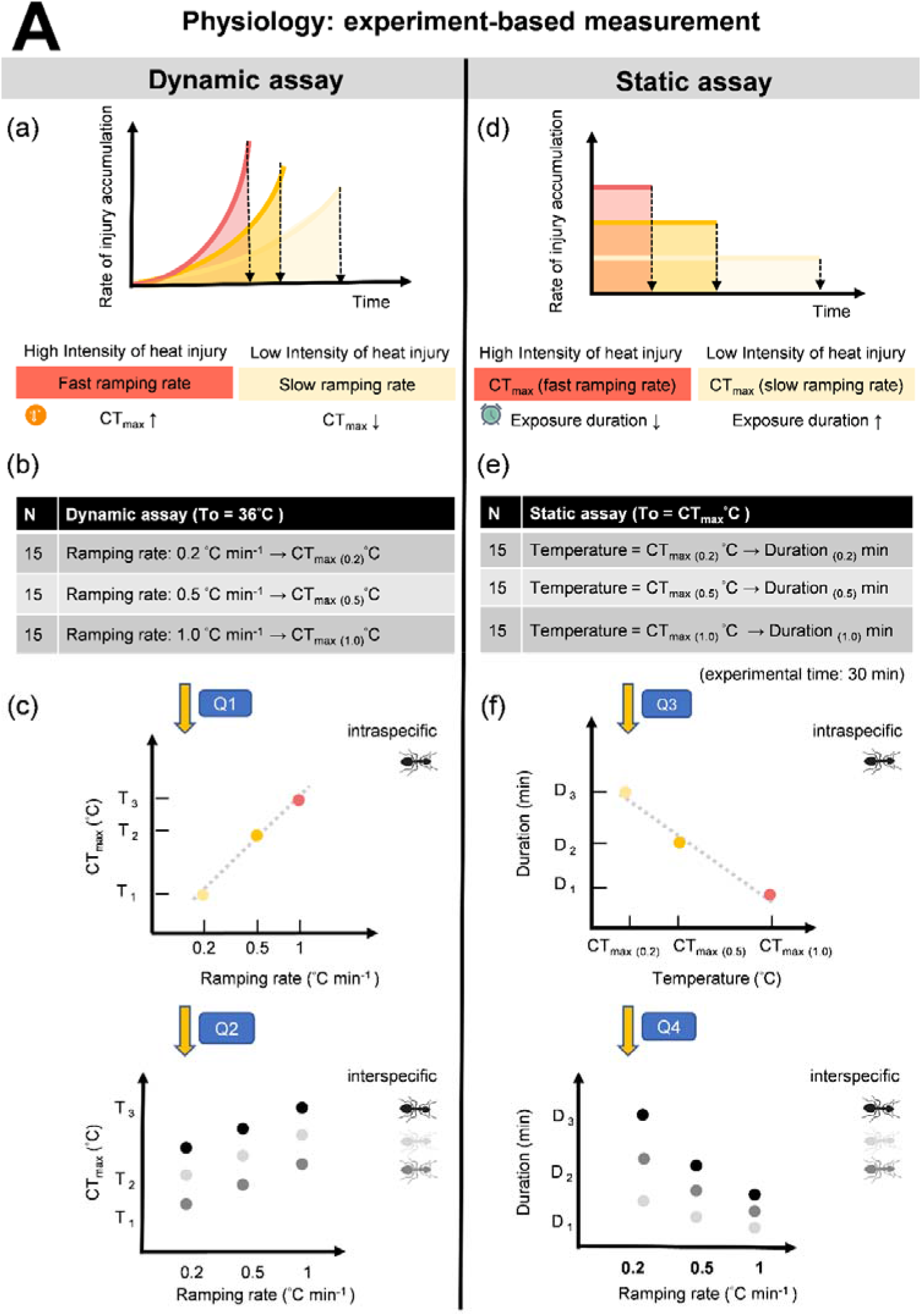
Experiment-based approach. (a) Theory of heat injury in dynamic assay for CT_max_ measurements: With a slow ramping rate (yellow curve), a longer exposure is needed to knock down the organism, than in an intermediate (orange curve) and a fast (red curve) ramping rate, because the injury accumulation rate increases slower with time. The knockdown time (inserted arrows) can be transformed to the knockdown temperature (CT_max_) using the ramp rate. (b) The experimental diagram of dynamic assay in this study with three treatments of ramping rate. (c) The aim of the first question (Q1) examines if a positive relationship between the ramping rate and CT_max_ values exists for each ant species at the intraspecific level, and (Q2) if a generalization of this relationship for all species is retrieved at the interspecific level. (d) Theory of heat injury in static assay for exposure duration: the injury accumulates at a temperature dependent constant rate during the static assay (yellow, orange, or red lines), and organism is knocked down once the species-specific amount of damage (time multiply temperature as the shaded area) has accumulated (e) The experimental diagram of static assay in this study with three temperature treatments (CT_max(0.2)_, CT_max(0.5)_ and CT_max(1.0)_. (f) The third question (Q3) examines if a negative relationship between temperature and duration exists for each ant species at the intraspecific level, and (Q4) interspecific level by measuring variation in CT_max_ values measured at three different ramping rates for 24 species.

**Figure 1B:**
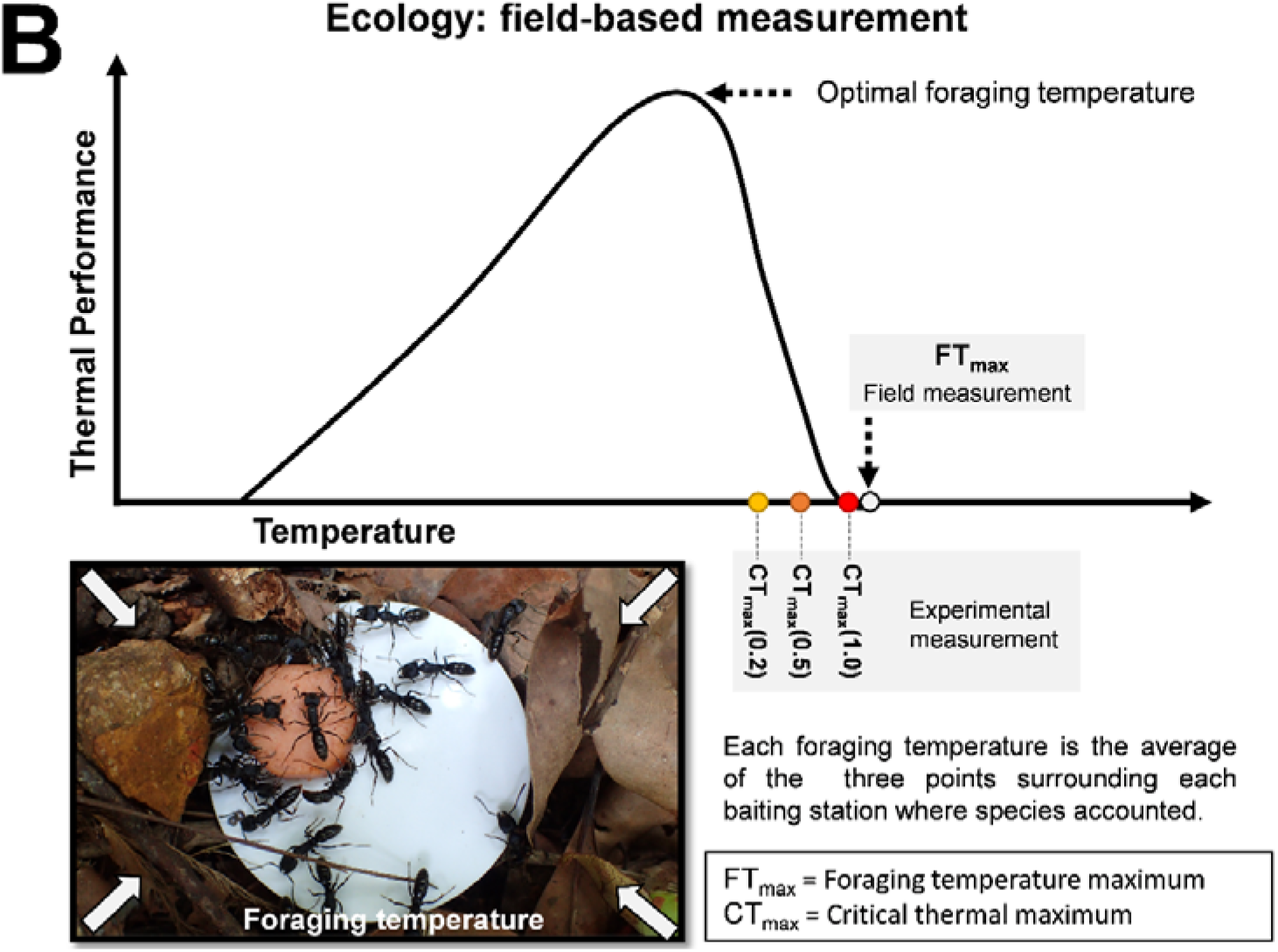
Field-based approach. Thermal performance curve of ectotherm on the basis of foraging behavior and the hypothesis between FT_max_ and CT_max_ retrieved by different ramping rates. Our study hypothesized that fast ramping rate can measure CT_max_ values in agreement with FT_max_ values. If specific ramping rate can measure the CT_max_ value present closest FT_max_, that ramping rate is more biologically relevant for CT_max_ to forecast the species activity within ecological studies. Methodology used to measure FT_max_ around baiting station is presented in the picture on lower left corner.

## 2. MATERIALS AND METHODS

### 2.1 Dynamic assay (Ramping temperatures to determine thermal limits)

To quantify upper thermal limit experimentally, dynamic assay of Critical Thermal maximum (CT_max_) used a continuous increase in temperature over time on organisms. Specifically, individuals were placed in an environment in which temperature was increased progressively and steadily according to a predefined ramping rate (°C per minute). In this study, general setting for measuring CT_max_ in ants was adapted from Bujan and Kaspari (2017) with three different ramping rates (see below) set as experimental treatments. We measured the CT_max_ of 27 species with different body size, microhabitat preferences on habitat and vertical stratum (see Supporting Information Section 2 for details of ant collection) using a digital dry bath (Benchmark - BSH1004, advertised accuracy ± 0.2°C). For each treatment, 15 individuals were tested, with a single individual ant worker placed within a 2.0 mL Eppendorf tube whose cap was filled with cotton to prevent individuals to hide into cooler position. In addition, to limit the stress experienced by individuals and the release of chemical defense (e.g. formic acid) which can be harmful in closed environment, individuals were led to the Eppendorf tube rather than being squeezed with forceps. Individuals were used only once. To ensure that the temperature indicated on the digital heat bath was accurate, an extra digital thermometer (UEi Test Instruments DT302 Dual Input IP67) inside a supplementary Eppendorf tube was placed as temperature control, and this temperature represents the most accurate experimental temperature of loss of muscle control.

Ramping rates were chosen based on a systematic review of previous studies (see Supporting Information Section 1 for details of literature collection), and also referred to environmental fluctuations observed within terrestrial ecosystems (Allen *et al.*, 2016; Stark *et al.*, 2017). Starting temperature was set at 36 °C without acclimation, which is the also lowest CT_max_ for all ant species known (Diamond *et al.*, 2012; Bennett *et al.*, 2018). Loss of muscle control was defined as an ending point for the individual, because it is more relevant to biological survival than lethal temperature (Lutterschmidt & Hutchison, 1997a). CT_max_ was recorded as the average temperature at which loss of muscle control occurred for each species at each ramping rate. Then, depending of the treatment tested, temperature was gradually increased by either 0.2, 0.5 or 1.0 °Cmin^−1^ (Fig. 1), until the individuals were observed having loss of muscle control (onset of spasm).

### 2.2 Static assay (duration in static heat temperature)

Static assay represents the experimental use of constant temperature to measure upper thermal limit until loss of muscle control in the certain dose of exposure duration. Individuals are placed in an environment with fixed temperature, with the duration during which they can tolerate such conditions is being recorded. Experimental temperatures used for static essays were retrieved from the CT_max_ values calculated from the previous dynamic experiment with ramping rates at 0.2, 0.5 or 1.0 °Cmin^−1^, such that for each species under three distinct static temperatures were tested (CT_max(0.2)_, CT_max(0.5)_, CT_max(1)_), and these temperatures being species specific (Fig. 1). The experimental setup and methods to record loss of muscle control were the same as in the dynamic Critical Thermal maximum assays, at the differences that temperature remained unchanged up to a maximum of 30 minutes. The checking time in dynamic and static assays were conducted every minute. However, if after 30 minutes the individuals did not lose muscle control, then the duration of those individuals in static assay was recorded as 30 minutes for data analyses.

### 2.3 Statistical analyses

#### 2.3.1 Dynamic and static assays

Before any analysis, individual CT_max_ was averaged by species identity and ramping rate. To assess if ramping rate significantly altered CT_max_ in dynamic and static assays, we conducted linear regressions, with CT_max_ values as the response and ramping rate as a continuous predictor. We used linear regression instead of ANOVA here, as the former provides adjusted R^2^, providing additional information on the performance of the model in explaining variations within the dataset. We also added species identity as another predictor to control for interspecific variations in CT_max_ and duration. We further conducted Tukey’s test to assess if increasing ramping rate leads to higher CT_max_ in dynamic assay and shorter duration in static assay in each species. To quantify the importance of ramping rate in determining CT_max_, we built a single predictor model with ramping rate as the only response, and reported the adjusted R^2^ values. For static assay, duration needed for loss of muscle control was log-transformed to normalize the data.

Different species experienced different variations of temperature. To examine if ramping rate affects comparison of species CT_max_ in 1) different strata, arboreal compared to ground and subterranean species and 2) urban relative to forest species, we built separate models, each with strata or habitat identity as the sole predictor. To examine whether the conclusions are sensitive to ramping rate, we built models for data obtained by different ramping rates separately. Thus, in total six models were built (three ramping rates x two factors). For the strata models, we further used Tukey’s test to assess if duration needed for loss of muscle control was higher in arboreal species. Ramping rate was treated as a continuous variable in all models. Additionally, for each species, we also assessed whether CT_max_ and duration needed for loss of muscle control significantly differed between the three ramping rates used. This was analyzed through treating ramping rate as a factor, and built linear models for each species, followed by Tukey’s test to conduct pairwise comparisons.

To minimize the potential impacts of variance heteroscedasticity, all linear regressions were adjusted using a heteroscedasticity-corrected coefficient covariance matrix (hc3) when assessing the significance of predictors (Long & Ervin, 2000). All statistical analyses were performed in R version 3.6.2 (R Development Core Team 2020). Linear models were built through function *lm*, followed by function *aov* in R-package *car* (Fox & Weisberg, 2018) to control for the effect of variance heteroscedasticity. Tukey’s post-hoc comparisons were done using *emmeans* (Lenth, Singmann & Love, 2018).

#### 2.3.2 Phylogenetic signal analyses

To test if ramping rate affects phylogenetic analysis, we used a backbone tree in genus-level phylogeny (Economo *et al.*, 2018), applying tree pruning to keep a single species for every genus in generating a genus-level phylogeny. Then we simulated 1,000 species-level phylogenies using Yule (pure-birth) process by function *genus.to.species.tree* in R-package phytools (Revell 2012). For each species-level phylogeny, we used *Pagel’s λ* and *Blomberg’s K* to examine phylogenetic signals in CT_max_ generated from different treatments. We also tested whether *λ* and *K* are significantly different from random using likelihood ratio test and randomization test (1,000 randomizations) respectively.

#### 2.3.3 Critical Thermal maximum vs. Foraging temperature maximum

For comparing CT_max_ and FT_max_ in each species (see Supporting Information Section 3 for details of foraging temperature collection), we used their absolute difference and calculated the mean and standard deviation for each ramping rate, as FT_max_ of each species was defined as the maximum foraging temperature observed across all the foraging temperature recorded.

## 3. RESULTS

### 3.1 Literature review of upper thermal limit in ant

A total of 52 publications (50 dynamic assay and 2 static assay studies) investigated ant upper thermal limits from 1944 to 2020 June (see the references used in the section of data availability). In total, 20 different ramping rate values were used, with 0.2 °Cmin^−1^ (13/50; 26%) and 1.0 °Cmin^−1^ (22/50; 44%) the most commonly used ramping rates (Fig. S1).

### 3.2 Ant species

A total of 2,934 individuals from 27 ant species representing eight subfamilies were used for dynamic and static assays, with three species, *Diacamma rugosum, Stigmatomma rothneyi*, and *Tetraponera microcarpa* omitted from static assay due to the insufficient number of individuals available.

### 3.3 Dynamic assay (Critical Thermal maximum CT_max_)

For all species, the CT_max_ values retrieved were dependent of the ramping rate used; with fast ramping rates produced significantly higher CT_max_ values than slow ramping rates (*p*-value < 0.001, Fig. 2, Table 1, Table S2). As a result, differences in the CT_max_ values retrieved between the slow ramping rate (0.2 °Cmin^−1^) and the fast ramping rate (1 °Cmin^−1^) averaged 4.13°C, ranging from 1.40 °C in *Aenictus* sp. *laeviceps* gp. to 6.47 °C in *Crematogaster rogenhoferi* (Table S2).

**Figure 2.**
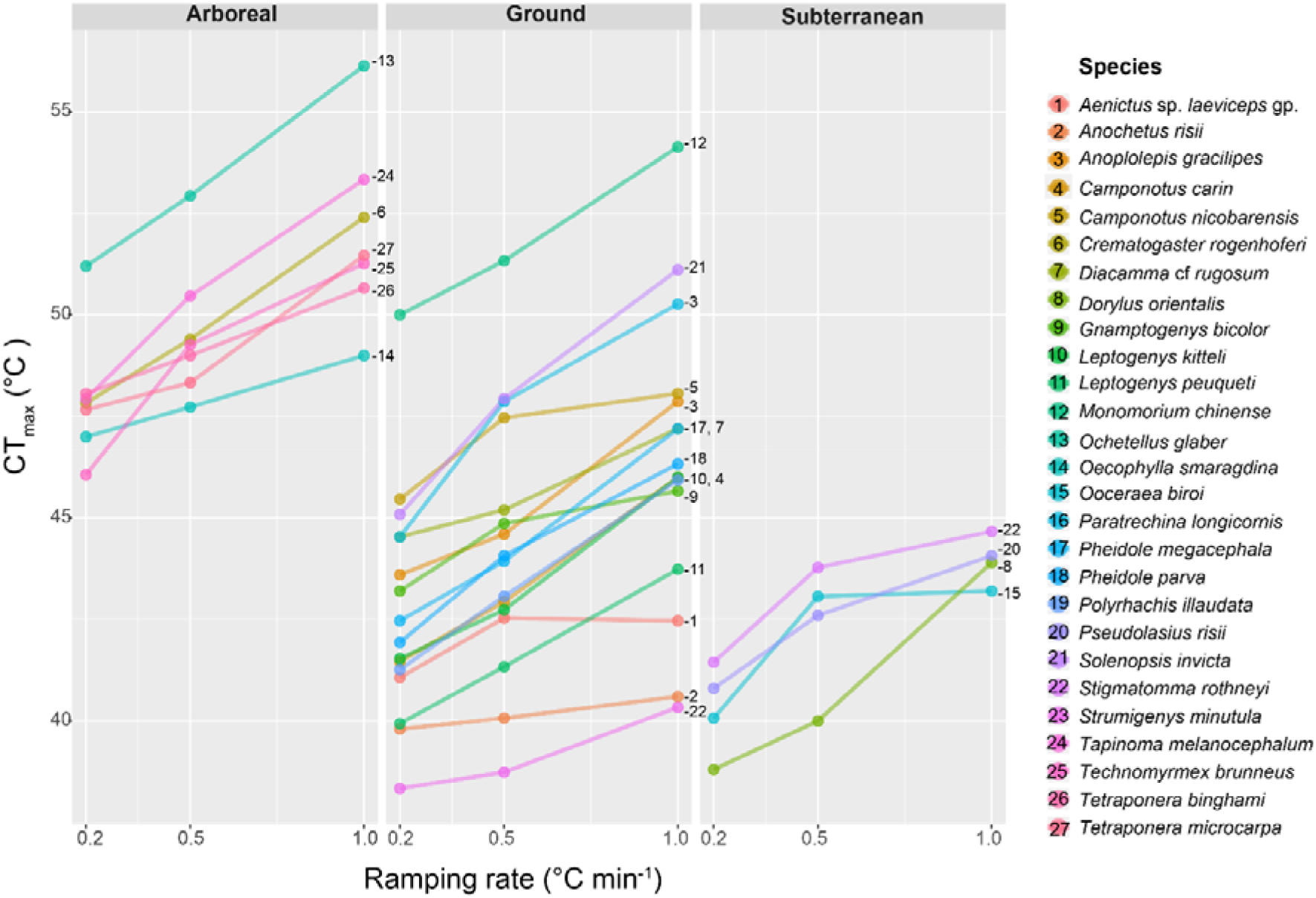
Dynamic assay: line plots of CT_max_ values measured in function of three ramping rates used for 27 ant species found in in function of their vertical stratification (arboreal, ground and subterranean strata).

**Table 1.** Outcomes of the linear models examining the relationship between parameters by heteroscedasticity-corrected covariance matrices linear model (1) dynamic assay: CT_max_, ramping rate and species identity (2) static assay: duration, ramping rate and species identity. We also reported the slope for ramping rate, as it is a continuous variable.

| Dynamic ($CT_{max}$) | Slope | df | F value | p-value |
| --- | --- | --- | --- | --- |
| Ramping rate | 4.7111 | 1 | 12.759 | <0.001 |
| **Adjusted R<sup>2</sup> = 0.1333** |  |  |  |  |
| Ramping rate | 4.7111 | 1 | 132.44 | <0.001 |
| Species identity | - | 26 | 235.30 | <0.001 |
| **Adjusted R<sup>2</sup> = 0.9344** |  |  |  |  |
| **Static (log<sub>10</sub>duration)** | **Slope** | **df** | **F value** | **p-value** |
| Ramping rate | -1.4949 | 1 | 90.444 | <0.001 |
| **Adjusted R<sup>2</sup> = 0.5078** |  |  |  |  |
| Ramping rate | -1.4949 | 1 | 97.496 | <0.001 |
| Species identity | - | 23 | 4.047 | <0.001 |
| **Adjusted R<sup>2</sup> = 0.7668** |  |  |  |  |

Generally, CT_max_ values for each species were correlated with the ramping rates used in the intraspecific model (Adjusted R^2^= 0.58 (0.11 - 0.94), *p*-value < 0.001; Table S4). When ramping rate was used as the only predictor in the interspecific model, CT_max_ values were positively correlated with the ramping rate (Adjusted R^2^= 0.13, *p*-value < 0.001; Table 1). Species identity was a stronger factor explaining CT_max_ values, as inclusion of species identity in the model strongly increased its explanatory power (Adjusted R^2^ for full model = 0.93, *p*-value for species identity < 0.001; Table 1).

Additionally, the CT_max_ values retrieved presented important differences in function of the ecology of the species. For instance, CT_max_ values for subterranean, ground and arboreal strata were significantly different for all three ramping rates (*p*-value < 0.001; Table S5), with higher thermal tolerance of arboreal species consistently higher than in ground and subterranean species for specific ramping rates (Post-Hoc test, *p*-value < 0.05). But CT_max_ values of species collected within urban habitats were not significantly different from species collected in forested habitats (*p*-value > 0.05) for all three ramping rates tested.

### 3.4 Static assay (exposure duration in static heat temperature)

Individual muscle control loss after 30 minutes was observed in all individuals of the 24 species tested for temperatures retrieved respectively for CT_max(0.5)_ and CT_max(1.0)_, but not for CT_max(0.2)_ for which four species had 40-96% of the 20 individuals tested presenting this condition (Table S3).

On average, increasing ramping rate significantly reduced the time needed for loss of muscle control in the interspecific model (*p*-value =<0.001 Table 1), with the duration for loss of muscle control at CT_max(0.2)_, CT_max(0.5)_, and CT_max(1.0)_ were 8.499 ± 4.295 min., 3.617 ± 1.507 min. and 2.244 ± 0.684 min. respectively (Fig. 3). For the majority of species, duration in response to loss of muscle control of each species under temperatures retrieved using different ramping rates were significantly different in the intraspecific model (*p*-value <0.05, Table S3). For the two extreme CT_max_ values (0.2 and 1.0 °Cmin^−1^) all species showed significant differences in their duration until muscle control loss (*p*-value < 0.05, Table S3).

**Figure 3.**
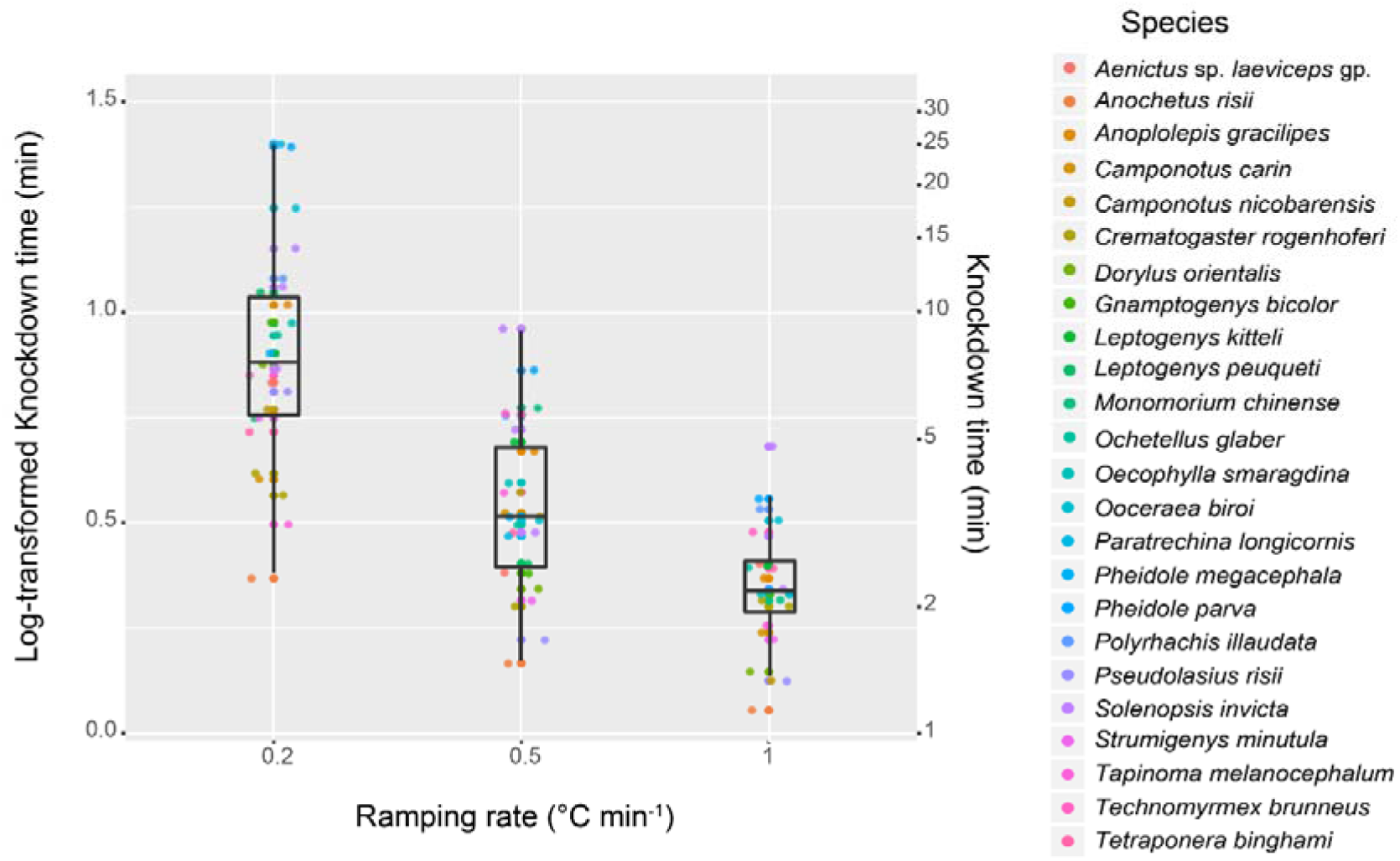
Static assay: Mean duration values (±SE) of 24 ant species for three temperatures based on the values retrieved in the CT_max(0.2, 0.5, and 1.0)_ assays.

Duration to muscle control loss was negatively correlated with ramping rate speed (Adjusted R^2^= 0.51, *p*-value < 0.001; Table 1). In addition to the ramping rate treatment used, species identity was an additional factor in determining the duration until muscle control loss (Adjusted R^2^= 0.77, *p*-value < 0.001; Table 1). Strata was not a significant factor in determining the duration to heat exposure that result in loss of muscle control; but habitat, i.e. urban and forest was a significant factor (*p*-value < 0.05; Table S5), with urban species presenting longer duration.

### 3.5 Phylogenetic signals

Methodological approaches used for measuring CT_max_ resulted in differences in the significance of phylogenetic signals. Differences were retrieved between 0.2, 0.5, and 1.0 °Cmin^−1^ in terms of how CT_max_ varied across ant phylogeny visually, with nine ant species (out of 27) showing obvious differences of phylogenetic signals (see: pink coloured species on Fig. 4). This was also confirmed through the use of the genus level polytomy tree and phylogenetic analyses, in which CT_max_ showed significant phylogenetic signal in both of *Pagel’s λ* (Mean = 0.991) and *Blomberg’s* K (Mean = 0.906) (Table S6) when measured at a ramping rate of 1.0 °Cmin^−1^ treatment, but lower values of *Pagel’s λ* (0.827-0.862) and *Blomberg’s* K (0.768-0.781) for the other two treatments (Table S6). Finally, the proportion of simulated trees that detected significant phylogenetic signals increased when higher ramping rate was used in measuring CT_max_.

**Figure 4.**
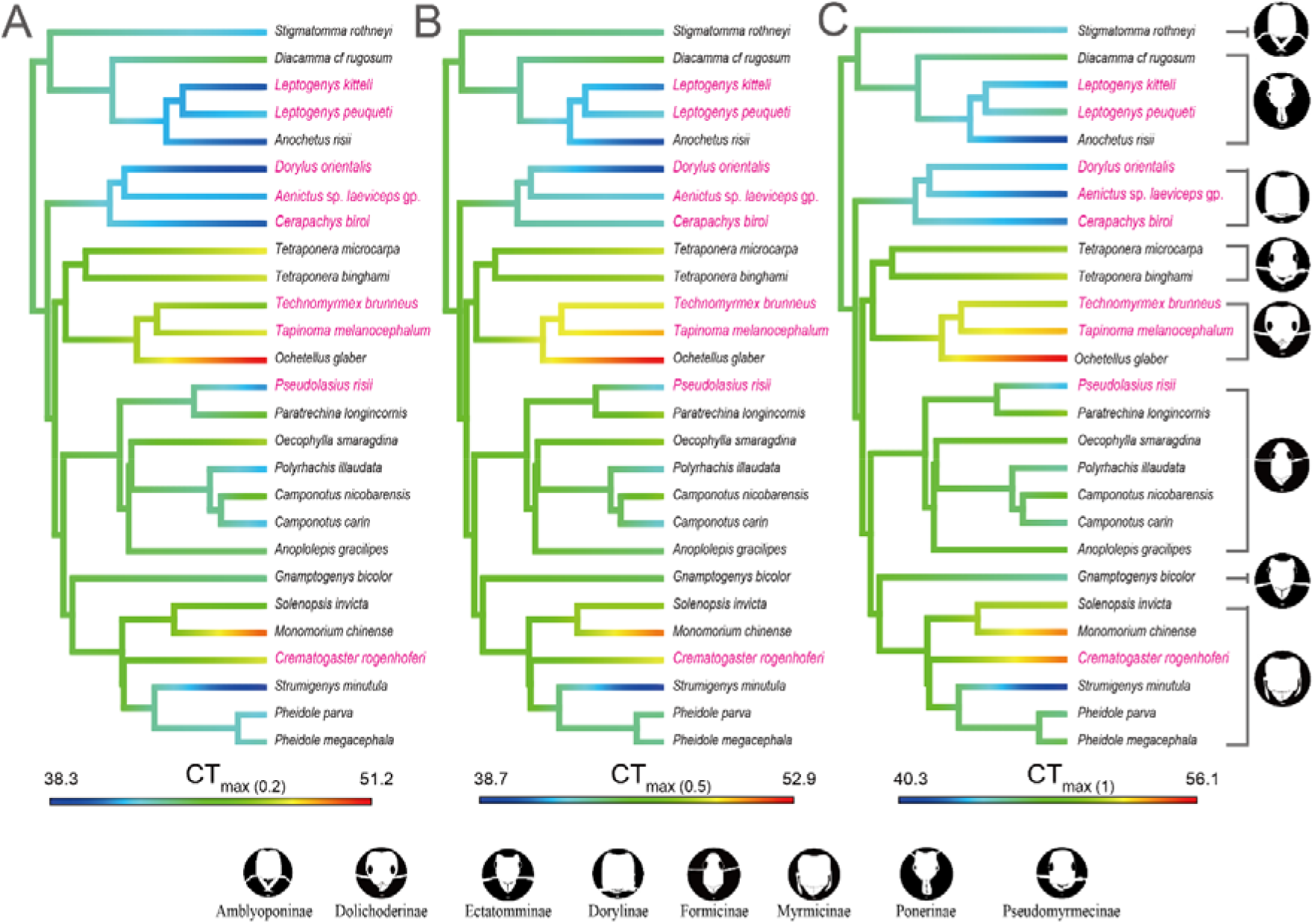
Critical Thermal maximum (CT_max_) of ant species in function of the phylogeny. Color shading corresponds with the magnitude of thermal tolerance measured with different ramping rates, A. 0.2 °Cmin^−1^, B. 0.5 °Cmin^−1^, C. 1.0 °Cmin^−1^. Ant illustrations credited to Mr Runxi Wang with permission.

### 3.6 Foraging temperature maximum vs. Critical Thermal maximum

The comparison of CT_max_ values retrieved from the three treatments show that CT_max(1.0)_ was the closest to the FT_max_ values measured for six out of seven species tested. Moreover, the absolute difference in FT_max_ and CT_max_ values retrieved show the best overall result for the 1 °Cmin^−1^ ramping rate (Mean ± SD: CT_max(1.0)_: 2.39 ± 1.41; CT_max(0.5)_: 3.79 ± 2.13; CT_max(0.2)_: 5.60 ± 2.57, Fig. 5 and Table S7), indicating a better overall performance in reconciling field and laboratory data.

**Figure 5.**
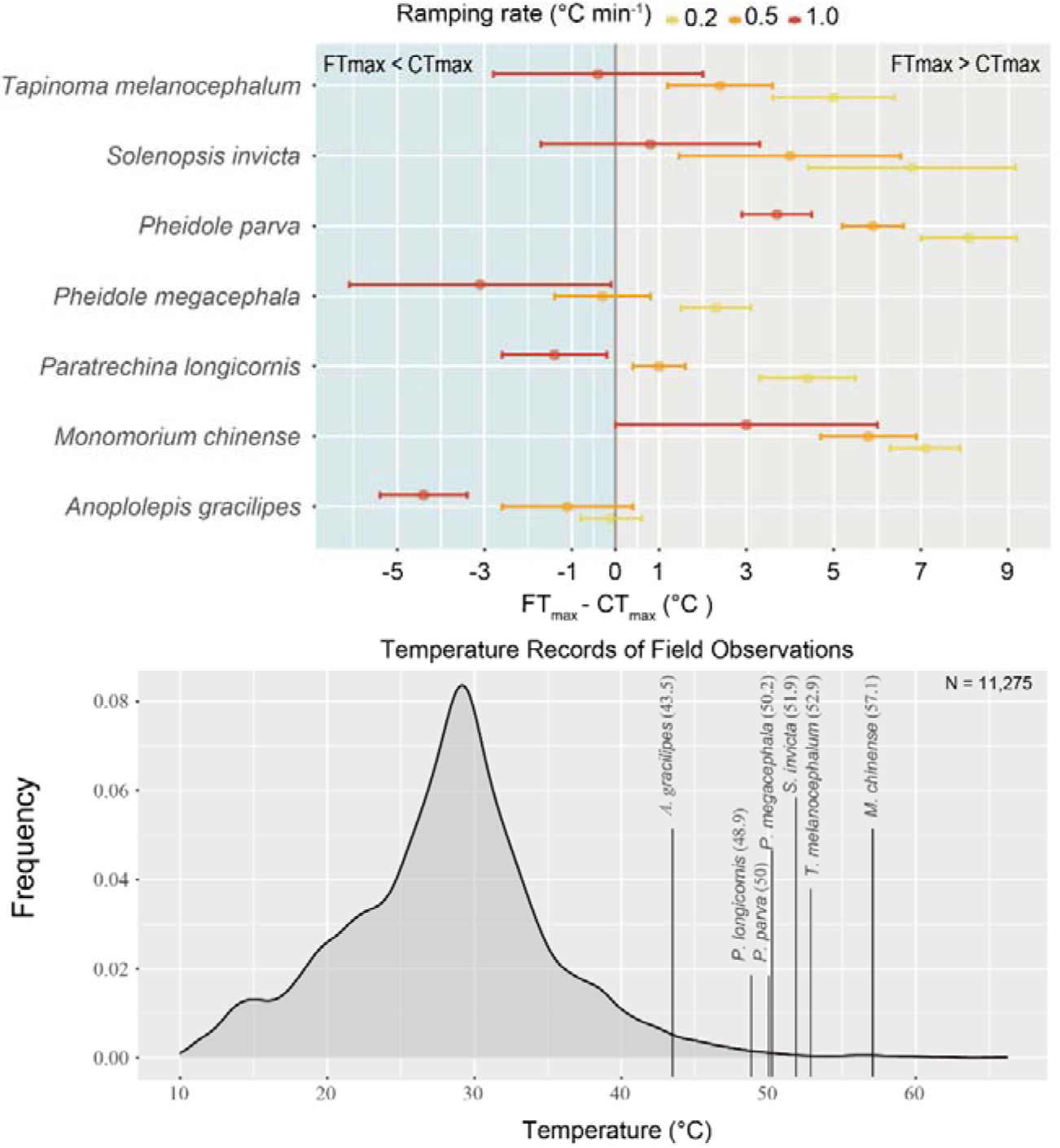
Upper plot showing the difference between FT_max_ and CT_max(0.2, 0.5, and 1.0)_ and errors bars as standard deviation of the CT_max_ values. Lower plot shows the range and distribution frequency of the surface temperatures measured near baiting stations during the sampling period, independently of the presence of ants or not. Vertical lines indicate the FT_max_ values measured for each species in the field.

## 4. DISCUSSION

Forecasting species warming tolerance using CT_max_ has received increasing attention (Lutterschmidt & Hutchison, 1997b; Bennett *et al.*, 2018). Although a satisfactory ramping rate has been discussed since the development of CT_max_, the use of slow and fast ramping is still debated with arguments based on ecological, physiological, and methodological aspects (Lutterschmidt & Hutchison, 1997b; Terblanche *et al.*, 2007; Chown *et al.*, 2009; Terblanche *et al.*, 2011; Jørgensen, Malte & Overgaard, 2019). Here, using a combination of dynamic and static assays on a wide range of species with distinct ecological, morphological, phylogenetic and biogeographic characteristics, our results show a consistent trend between ramping rate and CT_max_ values, and confirmed that an important part of variation observed in CT_max_ values results from the methodological approach used (Lutterschmidt & Hutchison, 1997b; Terblanche *et al.*, 2007; Jørgensen, Malte & Overgaard, 2019). We also demonstrated that the CT_max_ values retrieved from the fast ramping rate used (1°Cmin^−1^) not only represent a more realistic upper thermal limit in comparison to field data, but also influenced outcomes on phylogenetic analyses. Overall, we provided experimental and field-based support on designing a suitable ramping rate for ant CT_max_.

### 4.1 Literature review of upper thermal limit in ants

Dynamic and static assays represent two distinct approaches to measure thermal tolerance (Lutterschmidt & Hutchison, 1997b; Terblanche *et al.*, 2007). Our results show that in ants, an overwhelming number of studies (50/52) use the dynamic assay approach to assess upper thermal limits, confirming previous observations across a wide taxonomic range of studies (Lutterschmidt & Hutchison, 1997b). Our review also shows that a high variety of ramping rates (0.05 - 2.0 °Cmin^−1^) have been used to measure CT_max_ in ants, with a fivefold difference between the two most popular ramping rates e.g. 0.2 and 1.0 °Cmin^−1^ (Fig. S1). An emerging issue with such methodological differences is that results obtained from studies using different ramping rates are not directly comparable (Krebs, 1989); and unless corrections between CT_max_ and ramping rate are adopted, results from meta-analyses should be taken critically, with possible trends observed likely to be strongly impaired by methodological artifacts.

### 4.2 Ramping rate

The speed of ramping rate used in CT_max_ measurement is set either on ecological or physiological arguments. Ecology-based ramping rate suggests that ramping rate in CT_max_ assay should reflect the natural conditions or extreme events experienced by organisms (Terblanche *et al.*, 2011). Physiology-based ramping rate suggests that the ramping rate used in CT_max_ assay should induce individual’s body temperature to track the experimental ramping rate while limiting other side-effects such as thermal acclimation and longer exposure duration (Barker, Townsend & Hacunda, 1981; Hutchison & Murphy, 1985). As such, the ramping rate used in CT_max_ measurements is designed on the basis of physiological theory and approach, instead of being ecology-based.

Physiology-based ramping rate favors the use of fast ramping rate in measuring CT_max_ value (Barker, Townsend & Hacunda, 1981; Hutchison & Murphy, 1985). Specifically, the physiological theory and experiments support that 1 °Cmin^−1^, a rapid change in measuring CT_max_ values, provides valuable measures of the upper thermal limit. Although biophysics model claimed that ramping rates do not affect the synchronization between external temperature ***(Te)*** and individual body temperature ***(Tb)*** (Campbell & Norman 1998), thermal regulation inside body can have an effect on different ramping rates (Angilletta & Angilletta 2009). For example of fishes and frogs, fast ramping rate can allow the experimental ectotherm’s body temperature to follow the setting temperature and induce heat-shock effects independently of their size (Barker, Townsend & Hacunda, 1981; Hutchison & Murphy, 1985). However too fast or slow ramping rates show asynchronization in terms of inner body temperature (***Tb***) and experimental temperature (***Te***) (Barker, Townsend & Hacunda, 1981; Hutchison & Murphy, 1985). For instance, too fast ramping rates (e.g. 3.5, 3.8, and 10 °Cmin^−1^) cannot induce the heat-shock effects and cause delay effect (***Tb*** does not follow ***Te***); and too slow ramping rates (e.g. 1 °C hour^−1^, 5 °C hour^−1^, and 1 °C day^−1^) do not allow synchronization between ***Tb*** and ***Te*** and induce the thermal acclimation (Cox, 1974). Overall, fast ramping rate i.e. 1 °Cmin^−1^ used in CT_max_ is highly supported by physiological theories, as CT_max_ is ideally based on experimental approach measuring the physiological trait that represents species upper thermal limit (see: Cowles & Bogert, 1944; Hutchison & Murphy, 1985).

Ecology-based ramping rate was designed based on the environmental temperature variation of studying ectotherm, for example, researchers generally using faster ramping rate in terrestrial species than in aquatic species (Bennett *et al.*, 2018). The ramping rates, used by previous studies, were based on the species micro-habitat, thermo-equilibrium, etc. (Schumacher & Whitford, 1974; Lutterschmidt & Hutchison, 1997b; Terblanche *et al.*, 2007; Kaspari *et al.*, 2015). Some ecologists emphasized species’ micro-habitat and thus recommended that the measuring protocol should relate to the ectotherm thermal niches in order to compare species (Terblanche *et al.*, 2011). Nevertheless, our results do not support slow ramping rate for CT_max_ measurements, with these values failing to forecast species activity in function of ground temperature (see: foraging temperature section 3.6). Moreover, a majority of species show long tolerance duration (> 10 min.) in response to the CT_max_ values retrieved from slow ramping rate (see: static assay section 3.4 and Fig. 3), which thus does not represent a critical limit (Cowles & Bogert, 1944). It should be noted, however, that ecology-based ramping rate is still valuable in ecophysiology. Although ecology-based ramping does not reflect species’ upper thermal limit to withstand extreme temperature in the field (see the section of foraging temperature 4.3) and match the definition of CT_max_ (Cowles & Bogert, 1944), the ecology-based ramping may reflect thermal tolerance, and potentially species fitness, under specific temperature fluctuation (i.e. thermal niches, Connell, 1961; Lathlean, Seuront & Ng, 2017).

### 4.3 Foraging temperature maximum

Foraging temperature maximum (FT_max_) represents one of the most intuitive value derived from field observations and the identification of a maximum temperature threshold after which individuals of a species suspend their foraging activity. Although FT_max_ represents upper thermal threshold of species activity in the field (Whitford & Ettershank, 1975), recording of FT_max_ also includes several uncontrolled and cofounding factors such as weather conditions and species interactions, which might ultimately affect the values measured (Bruno, Stachowicz & Bertness, 2003). Alternatively, the use of CT_max_ can be used independently of the habitat conditions to estimate upper thermal limits. If the measurement of CT_max_, is relatively easier than FT_max_, questioning its biological relevance in relation to the methodology used is paramount (Lutterschmidt & Hutchison, 1997b). Here, we attempted to identify the most suitable ramping rate in CT_max_ measurements through a comparison with FT_max_ values. Because CT_max_ is supposed to represent a maximum physiological threshold, one should thus expect for those values to top FT_max_ values at which individuals are actively foraging. If CT_max_ values are inferior to FT_max_ values, then those, theoretically, indicate a methodological underestimation. Our result shows discrepancies between FT_max_ and CT_max_ values, which are particularly marked for the slowest ramping rate (0.2 °Cmin^−1^) for six species (out of 7) with CT_max_ values 4.4 °C to 8.1 °C inferior to the FT_max_ retrieved. Such gaps question the biological relevance of slow ramping rate CT_max_ values. In contrast, a majority of CT_max_ values retrieved with highest ramping rate (1 °Cmin^−1^) aligned more precisely with the FT_max_ values retrieved (Fig. 5 and Table S7).

Temperature is one of the most important factors to predict species distribution, in particular for ectotherms, which activity is limited by ambient temperature (Araújo *et al.*, 2013). FT_max_ and CT_max_ are widely used in Species Distribution Model, and a recent study shown that FT_max_ provides more accurate modelling predictions than CT_max_ (Guo *et al.*, 2020). Although FT_max_ is advantageous and precise in forecasting species activity and distribution, its measurement is challenging. To determine FT_max_, researchers used microhabitat temperature such as surface temperature and thus represent a more realistic measurements of the conditions experienced by species rather than air temperature (Kaspari & Weiser, 2007; Spicer *et al.*, 2017) and some of the confounding effects mentioned above. Moreover, the habitat in which a species is most commonly encountered might not always experience temperature close to their thermal limit, such as for species living in the leaf-litter layer of closed-canopy forests. For instance, in our results, the lowest CT_max_ value observed, independently of the ramping rate used is of 38.3 °C, which remains 0.5 °C superior to the highest air temperature on record within Hong Kong (Hong Kong observatory), and in the absence of direct solar radiations, soil temperatures are similar or inferior to air temperature (Alam *et al.*, 2015). Thus, for some habitats or microhabitats, temperatures measured in the field cannot be translated directly to a maximum temperature threshold. Therefore, the measurements of CT_max_ based on experimental approach remains necessary as long as it can be standardized and represent biologically relevant threshold.

Our results show that within a species, the use of a particular ramping rate could lead to ~1.5 to 6.5 °C differences in CT_max_ measurements, which would certainly lead to tremendous differences in species distribution models and forecasts in responses of climate change. Although the speed of ramping rate does not affect thermal tolerance comparison on the ant species from strata/habitats (Fig. S5), the CT_max_ retrieved can affect species distribution modeling (Guo *et al.*, 2020) and forecast of species activity with coupling environmental temperature. Biologically relevant CT_max_ is paramount to forecast species activity in function of temperature, while biologically inaccurate CT_max_ mischaracterize species activity and distribution. The Red Imported Fire Ant example, *(Solenopsis invicta)* with populations in Hong Kong introduced from the USA (Ascunce *et al.*, 2011), is relevant to illustrate this,. The CT_max_ of this species in the USA has been measured using slow ramping rate of 0.12 or 0.2 °Cmin^−1^ (Bentley, Hahn & Oi, 2016; Roeder, Roeder & Kaspari, 2018) or faster ramping rate 1.0 °Cmin^−1^ (Bentley, Hahn & Oi, 2016; Wendt & Verble-Pearson, 2016). Coincidently, our field observations on the fire ant activity (N = 1,398) corresponds to the thermal threshold observed for the fast ramping rate in both the USA and in Hong Kong (this study), with a difference of 0.71 °C on average, but not with those retrieved at slow ramping rate which are superior by 4.77 °C on average (FT_max_> CT_max_). While few intensive activity-temperature studies of ectotherms like those on *S. invicta* are available (Vogt *et al.*,2003; Drees, Summerlin & Vinson, 2007; Roeder, Roeder & Kaspari, 2018), those field observations showed the importance of biologically relevant CT_max_ values to predict species activity, not only for single species but also entire species assemblage (see: section 3.6 and Fig. 5).

### 4.4 Phylogenetic signals

Ectotherm heat tolerance exhibited strong phylogenetic conservatism in most studies, including ants (Diamond & Chick, 2018; Bujan *et al.*, 2020), fruit flies (Kellermann *et al.*,2012a), or lizards (Grigg & Buckley, 2013). In contrast, other studies retrieved no evidence for a relationship between phylogeny and thermal tolerance (Bishop *et al.*, 2017; Nowakowski *et al.*, 2018). To the best of our knowledge, our results represent the first comparative study between phylogenetic signal and ramping rates and show that the detection of a phylogenetic signal (Pagel’s *λ* and Bloomberg’s *K)* is directly influenced by the methodology used to measure CT_max_ (Table S6). It also shows that apart from differences in tree topography and species pool, differences in ramping rate could also explain these inconsistencies. Overall, heat tolerance has been shown to be strongly constrained by evolutionary history (Diamond & Chick, 2018), the consideration of the ramping rates used to collect data should also be integrated as an important cofounding factor for testing evolutionary hypotheses. To produce such analyses, the use of slow ramping rate or the mix of values originating from different methodologies should be avoided.

### 4.5 Conclusion

The use of CT_max_ to study ectotherms has significantly increased over the past century, and its application has yielded multiple predictions about global change impacts (Stillman, 2003; Kellermann & van Heerwaarden, 2019). Our results show that CT_max_ can represent the upper thermal limits of ants and be biologically significant but needs to be measured through the proper ramping rate. The proper ramping rate to measure CT_max_ should be designed on the basis of taxon-group (Kovacevic, Latombe & Chown, 2019). Although the present ant studies and *Drosophila* studies present similar correlation between CT_max_ and ramping rate in the static and dynamic assays (Jørgensen, Malte & Overgaard, 2019; Rezende *et al.*, 2020), we found that CT_max_ measurements through the “best” ramping rate can be a reliable predictor for ant foraging activity and associated maximum temperatures measured in the field. This comparison suggests that ramping rate used in CT_max_ measurements should also be tested against species upper thermal limit measured in the field (e.g. FT_max_). Our study thus established a new, hybrid method, integrating dynamic and static assays and comparison with FT_max_, to seek a suitable ramping rate, and we advocate for the standardization of 1 °Cmin^−1^ to measure ant CT_max_ values. This novel integrative approach should be extended to other ectotherms to identify suitable ramping rate. The accurate measurement of CT_max_ holds the potential to reveal crucial information of species upper thermal limit, with important implications for climate warming, land-use change, biodiversity conservation, pest control and trait-based research.

## ACKNOWLEDGMENTS

We are grateful to the HKU IBBL members, Runxi Wang (illustrations on Fig. 4), Mark Wong, Francois Brassard, Roy Cheung, Roger Lee, Kin Chan, Mac Pierce, Brian Worthington and Maria Lo for their help and suggestions; Timothy Bonebrake, Dr Lisa Bjerregaard Jørgensen, Dr Jelena Bujan, Dr Bay den D. Russell for valuable suggestions. CML is supported by Fundação Macau and Direcção dos Serviços do Ensino Superior. CML, TPNT, and BG are supported by The University of Hong Kong and an Early Career Scheme 2017/18 grant (27106417).

## AUTHORS’ CONTRIBUTIONS

According to the Contributor Roles Taxonomy (CRediT) framework, in this study: Conceptualization by **CML** and **BG**; Funding acquisition by **CML** and **BG**; Supervision by **BG**; Investigation by **CML**; Methodology by **CML** and **BG**; Data curation by **CML**; Formal analysis by **CML** and **TPNT**; Writing-original draft by **CML**; Writing-review & editing by all authors.

## DATA AVAILABILITY

Data later will be available from the Dryad Digital Repository

